# *Fusobacterium nucleatum* facilitates cetuximab resistance in colorectal cancer via the PI3K/AKT and JAK/STAT3 pathways

**DOI:** 10.1101/2020.12.21.423755

**Authors:** Hu Han, Yan Li, Wan Qin, Lu Wang, Han Yin, Beibei Su, Xianglin Yuan

**Affiliations:** Department of Oncology, Tongji Hospital, Huazhong University of Science and Technology, Wuhan, Hubei Province, China; Department of Oncology, First Affiliated Hospital, School of Medicine, Shihezi University, Shihezi, Xinjiang Province, China; Medical Department, First Affiliated Hospital, School of Medicine, Shihezi University, Shihezi, Xinjiang Province, China

**Keywords:** *Fusobacterium nucleatum*, colorectal cancer, cetuximab, resistance, PI3K/AKT, JAK/STAT3

## Abstract

Infectious pathogens contribute to about 20% of the total tumor burden. *Fusobacterium nucleatum* (*Fn*) has been associated with the initiation, progression, and therapy resistance in colorectal cancer (CRC). The over-abundance of *Fn* has been observed in patients with right-sided CRC than in those with left-sided CRC. While the *KRAS/NRAS/BRAF* wild-type status of the CRC conferred better response to cetuximab in patients with left-sided CRC than with right-sided CRC. However, treatment failure remains the leading cause of tumor relapse and poor clinical outcome in patients with CRC. Here, we have studied the association of *Fn* to cetuximab resistance. Our functional studies indicate that *Fn* facilitates resistance of CRC to cetuximab *in vitro* and *in vivo*. Moreover, *Fn* was found to target the PI3K/AKT and JAK/STAT3 pathways, which altered the response to cetuximab therapy. Therefore, assessing the levels and targeting *Fn* and the associated signaling pathways may allow modulating the treatment regimen and improve prognoses of CRC patients.

## Introduction

CRC accounts for a high incidence and mortality rate, globally^1^, and arises due to complex interactions of genetic attributes, lifestyle, and environmental factors^2^. The treatment objectives in patients with metastatic CRC (mCRC) include, reduction of cancer load, cancer growth inhibition, prolonging survival time, and improving the quality of life. Monoclonal antibodies (mAbs) against the epidermal growth factor receptor (EGFR), such as cetuximab and panitumumab, bind the extracellular structural domain of the EGFR, and reinforce receptor intemalization and degeneration^3,4^. These anti-EGFR mAbs have been considered as targeted therapy for the treatment of mCRC patients harboring *KRAS* wild type (WT) status^5,6^. Furthermore, when administered as a monotherapy, cetuximab may confer persistent responses in 12-17% of the CRC patients^7^, and provide about 72% reaction rate as a combined chemotherapy^8^.

However, several clinical trials have indicated that patients with left-sided mCRC harboring *RAS/BRAF* WT status showed better response to cetuximab than those with right-sided mCRC^7–19^. Moreover, clinical data indicates that the response achieved in CRC with *RAS/BRAF* WT status is transient, in the best responders show recurrence within 18 months of treatment^20^. Furthermore, patients frequently present with chemoresistance, and only 10-20% mCRC patients benefit from the mAb treatment^20,21^. Several studies have described the mechanisms for intrinsic and acquired resistance to treatment with anti-EGFR mAbs, which we have reviewed previously^22^. In brief, these mechanisms mainly include mutations in the *RAS, BRAF, PIK3CA*, and *EGFR*, and phosphorylation of the STAT3 and AKT, which induce resistance to the treatment with anti-EGFR mAbs, primarily via persistent activation of the EGFR downstream signaling pathways despite the EGFR blockade. Deregulation of these PI3K/AKT and JAK/STAT3 signaling pathways inhibit apoptosis, and promote multiple drug resistant in tumor cells^23,24^ However, the non-genetic and/or non-epigenetic mechanisms of resistance remain to be explored in details. Further, drug resistance leads to tumor recurrence, and the 5-year survival rate in unresectable mCRC patients remains less than 10%^25^. Additionally, patients rarely respond to immunosuppressive agents^26^. Therefore, it is necessary to understand the mechanism of resistance to cetuximab in patients with CRC.

Recent reports have indicated the gut microorganisms to be associated with CRC^27–31^. Further, several studies have validated that tumor tissues and fecal specimens of CRC patients show high amounts of *Fn* than in normal controls^32–37^ Furthermore, the abundance of *Fn* was observed in patients with right-sided CRC than in those with leftsided CRC^38–40^. Moreover, CRC associated with *Fn* presents with worse clinical outcome^36,41,42^. Additionally, *Fn* enhances tumor growth^43^, protects tumors from immune attack^44^, and promotes chemo-resistance^42^. Taken together, *Fn* appears to be not only enriched in CRC but also to promote tumor initiation, growth, progression, and therapy resistance in patients.

To understand how cetuximab confers varied response in patients with right-sided and left-sided CRC, we hypothesized that the presence of *Fn* may contribute to therapy resistance. A thorough literature survey indicated that the role of *Fn* in resistance to anti-EGFR mAbs remains to be explored in human CRC. Therefore, here, we have explored the association of *Fn* with resistance to cetuximab using an *in vitro* and *in vivo* approach. Our analysis implicates the PI3K/AKT and JAK/STAT3 pathways to confer resistance to cetuximab in response to *Fn* in CRC cells. Moreover, given the high proportion of cetuximab resistance in CRC patients, identification of such amenable targets should allow designing better treatment modalities and improve the survival outcome in patients.

## Materials and Methods

### Bacterial strain and cultured conditions

*Fusobacterium nucleatum* strain ATCC 25586^32^, were purchased from China General Microbial Species Preservation Center (CGMCC). *Fn* was cultured overnight under anaerobic conditions at 37 °C in brain heart infusion broth supplemented with Vitamin K1, hemin, and L-Cysteine as described before^45^.

### Cell lines and drug

CRC cell lines SW620, SW480, SW48, Caco2, LOVO, HT29, and HCT116 were cultured in DMEM medium (GIBCO, Carlsbad, CA) supplemented with 10% fetal bovine serum (FBS) (Hyclone; GE Healthcare Life Sciences) at 37°C in a humidified 5% CO_2_ atmosphere. All CRC cell lines were tested to exclude mycoplasma contamination. Cetuximab (5 mg/mL) was kindly provided by Merck (Darmstadt, Germany).

### Mutation analysis

DNA isolation from the CRC cell lines SW620, SW480, SW48, Caco2, LOVO, HT29, and HCT116 were performed making use of the QiAmp DNA-Isolation Kit (Qiagen, Hilden, Germany) in accordance with the instructions. *KRAS* and *PIK3CA* polymerase chain reactions for *KRAS* exon-2,3 and *PIK3CA* exon-9,20 augmentation were managed employing the primers listed in the table (See supplementary materials). The products of PCR were submited to direct sequencing. The results of sequencing were compared to the *KRAS* and *PIK3CA* sequence stored in GenBank databases.

### Cell viability assay

The viability assays were determined in Caco2 and HT29 cells. CRC cells were seeded in 96-well plates at 2×10^3^ cells per well with 100 μL culture medium.

The first approach: 24 h after seeding, cells were treated with *Fn* at various multiplicity of infection (MOI) of 100:1, 500:1, 1000:1. Cells were treated with medium as a negative control. Cell absorbance was detected by spectrophotometer in Caco2 and HT29 cells at indicated time.

The second approach: 24 h after seeding, cells were treated with cetuximab (0.1, 1, 10, 100, 1000 μg/ml) with or without *Fn* at MOI (100:1) for 48h. Cells were treated with medium as a negative control. Cell absorbance was detected by spectrophotometer in Caco2 and HT29 cells at indicated time.

After indicated time, cell viability was analysed employing a Cell Counting Kit in line with the manufacturer’s instructions (CCK-8; Dojindo, Kumamoto, Japan). The half maximal inhibitory concentration (IC50) values were computed employing Graph-Pad Prism 6.0 (GraphPad Software, San Diego, CA, USA). According to the results, we chose optimal dose of *Fn* and concentration (cetuximab) to continue our experiments. Each experiment was repeated three times.

### Cell proliferation assay

Proliferation assays were determined in Caco2 and HT29 cells. 1×10^5^ cells were seeded in per well of 6-well plates supplemented with 4 ml culture medium per well. The medium was replaced the second day. Then, cells were treated with cetuximab (100μg/ml) and/or *Fn* at MOI 100:1. Cells were treated with medium as a negative control. The numbers of cell were counted in indicated time using a hemocytometer. Each experiment was repeated three times.

### Colony formation assay

Cells were seeded at 500 cells per well in six-well plates. 24h incubation, and then the cells treated with cetuximab (100 μg/ml) and/or *Fn* at MOI (100:1), and then continuously incubated in fresh medium. Cells were treated with medium as a negative control. After 14-days incubation, the cells were washed twice with phosphate buffer solution, fixed for 20 min with 10% paraformaldehyde, and stained for 20 min with 0.05% crystal violet. The visual colonies were calculated.

### Cell Cycle Analysis

Caco2 and HT29 cells were seeded at a density of 1×10^5^ cells per well in 6-well plates. 24 h incubation, and then cells were treated with the same manipulation as mentioned (colony formation assay) for 48h. The adherent fractions of cells were trypsinized and fixed overnight with 100% ice-cold ethanol at −20 °C. The cells were washed twice with cold PBS and stained with RNase a and PI solutions in the dark at 37 °C for 30 minutes. The cell cycle distribution was analyzed by flow cytometry (BD Biosciences, San Jose, CA). Each experiment was repeated three times.

### Cell apoptosis

Caco2 and HT29 cells were seeded at a density of 1×10^5^ cells per well in 6-well plates. 24 h incubation, and then the cells were treated with cetuximab (100μg/ml) and/or *Fn* at MOI 100:1. Cells were treated with medium as a negative control. The extent of apoptosis was detected via Annexin V-FITC/PI apoptosis detection kit in line with the manufacturer’s instruction. The cells were immediately analyzed by flow cytometry (BD Biosciences, San Jose, CA). Each experiment was repeated three times.

### Tumor Xenograft Study

Four-week-old male athymic nude mice were raised under specific pathogen-free conditions provided with food and water provided ad will. Each mouse was subcutaneously injected with 1×10^7^ Caco2 cells in the right axilla to set up a xenograft model. After 7 days of inoculation, the mice were randomly divided into 4 groups. There were four groups: i) saline (Control); ii) *Fn* bacteria solution; iii) cetuximab (1 mg/mice); iv) cetuximab (1 mg/mice) and *Fn* bacteria solution. *Fn* (1×10^7^ Clone forming unit (CFU)) was given by multipoint intratumoral injection^42^, twice a week for three weeks. Cetuximab treatment was given at a dose of 1 mg/mouse, intraperitoneal injection, twice a week for three weeks^46^.

Tumor volume and weight were calculated performed as previously described^42^. The research procedures were approved by the Institutional Animal Care and Use Committee of Tongji Hospital, Tongji Medicine, Huazhong University of Science and Technology.

### Western Blot Assays

Total cell proteins were extracted after treatment with medium (control) and *Fn* at appointed time. The assays were carried out as previously described^47^. The target proteins were detected with primary antibodies recognizing AKT (Cell Signaling Technology, CST #9272; 1:1,000), p-AKT (CST #4060; 1:2,000), STAT3 (CST #9139; 1:1,000), and p-STAT3 (CST #9145; 1:2,000). GAPDH (CST #5174; 1:1,000) was used as a loading control.

### Statistical Analysis

All results are expressed as means ± standard errors of means (SEM) of three independent experiments. Statistical analysis was carried out employing Graph-Pad Prism 6.0 (GraphPad Software, SanDiego, CA, USA). Statistical significance was reported if the *p* value was <0.05 employing an unpaired Student’s *t*-test.

### Statement

The research procedures were approved by the Institutional Animal Care and Use Committee of Tongji Hospital, Tongji Medicine, Huazhong University of Science and Technology, including any relevant details; We confirmed that all experiments were performed in accordance with the relevant guidelines and regulations.

## Results

### Selection of multiplicity of infection (MOI) and IC50

To understand whether *Fn* participates in resistance to cetuximab, *Fn* was co-cultured with Caco2 and HT29 CRC cell lines using varied MOI. Our analysis suggested that *Fn* (MOI = 100) did not affect the proliferation of these cells (Figure 1A and 1B).

**Figure 1.**
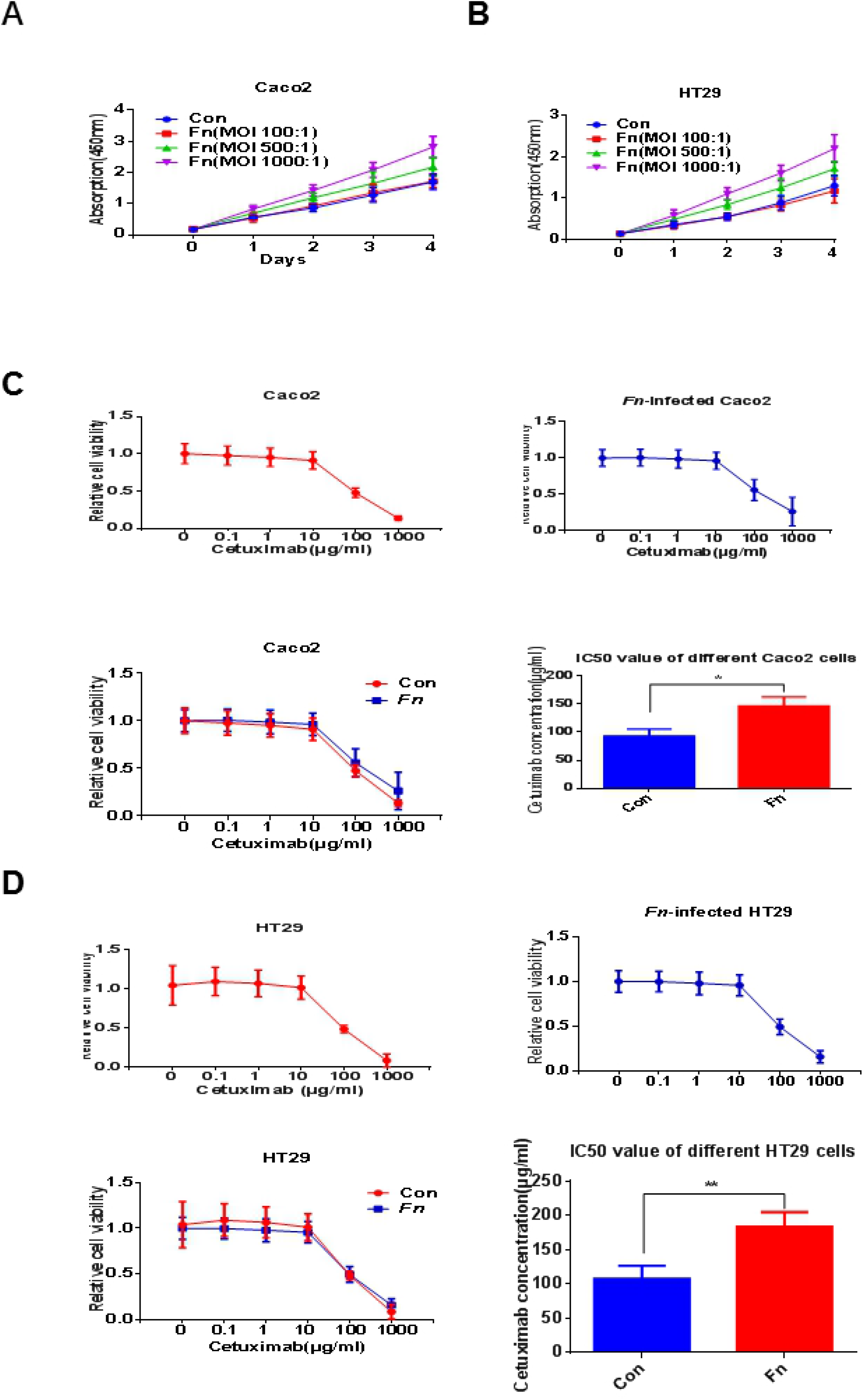
Selection of multiplicity of infection (MOI) and IC50. (A and B) Cell absorption was measured by spectrophotometer in Caco2 and HT29 cells co-cultured with *Fn* at different MOIs. All analyses were performed using unpaired Student *t* test. (C) Cell viability was detected by CCK-8 in parental Caco2 cells and parental Caco2 cells co-cultured with *Fn* (MOI:1OO) in the presence of different concentrations of cetuximab. All analyses were performed using unpaired Student *t* test. (D) Cell viability was detected by CCK-8 in parental HT29 cells and parental HT29 cells co-cultured with *Fn* (MOI:1OO) in the presence of different concentrations of cetuximab. All analyses were performed using unpaired Student *t* test. \**P* < 0.05, \*\**P* < 0.01, \*\*\**P* < 0.001 by unpaired Student *t* test. Con, control; *Fn, Fusobacterium nucleatum*; Cet, cetuximab.

To quantitatively assess the effect of *Fn* on resistance to cetuximab, we compared cetuximab-induced cell viability among the parental Caco2 and HT29 cells, and the *Fn*-co-cultured Caco2 and HT29 cells, in the presence of different concentrations of cetuximab. Our analysis suggested the half maximal inhibitory concentration (IC50) to be 93.47 ± 6.978 μg/ml and 146.4 ± 9.430 μg/ml in the parental Caco2 cells and *Fn*-co-cultured Caco2 cells, respectively (Figure 1C *p*=0.0107). Whereas, the IC50 for parental HT29 cells and the *Fn*-co-cultured parental HT29 cells was 107.8 ± 10.91 μg/ml and 183.7 ± 12.28 μg/ml in the, respectively (Figure 1D *p* = 0.0099). Moreover, these differences in IC50 values for parental and *Fn*-co-cultured cells were statistically significant, indicating that *Fn* (MOI= 100) conferred cetuximab resistance in the Caco2 and HT29 CRC cells.

### *Fn* induces cetuximab resistance in CRC cells *in vitro* and *in vivo*

While the Caco2 and HT29 cells are relatively sensitive to cetuximab treatment^46,48,49^, we confirmed their *KRAS/PIK3CA* (WT) status. Consistent with our hypothesis, cetuximab inhibited cell proliferation in parental Caco2 and HT29 cells but not in cells co-cultured with *Fn* (Figure 2A and 2B). Therefore, these data indicate that *Fn* inhibits the effects of cetuximab on cell proliferation in Caco2 and HT29 cells.

**Figure 2.**
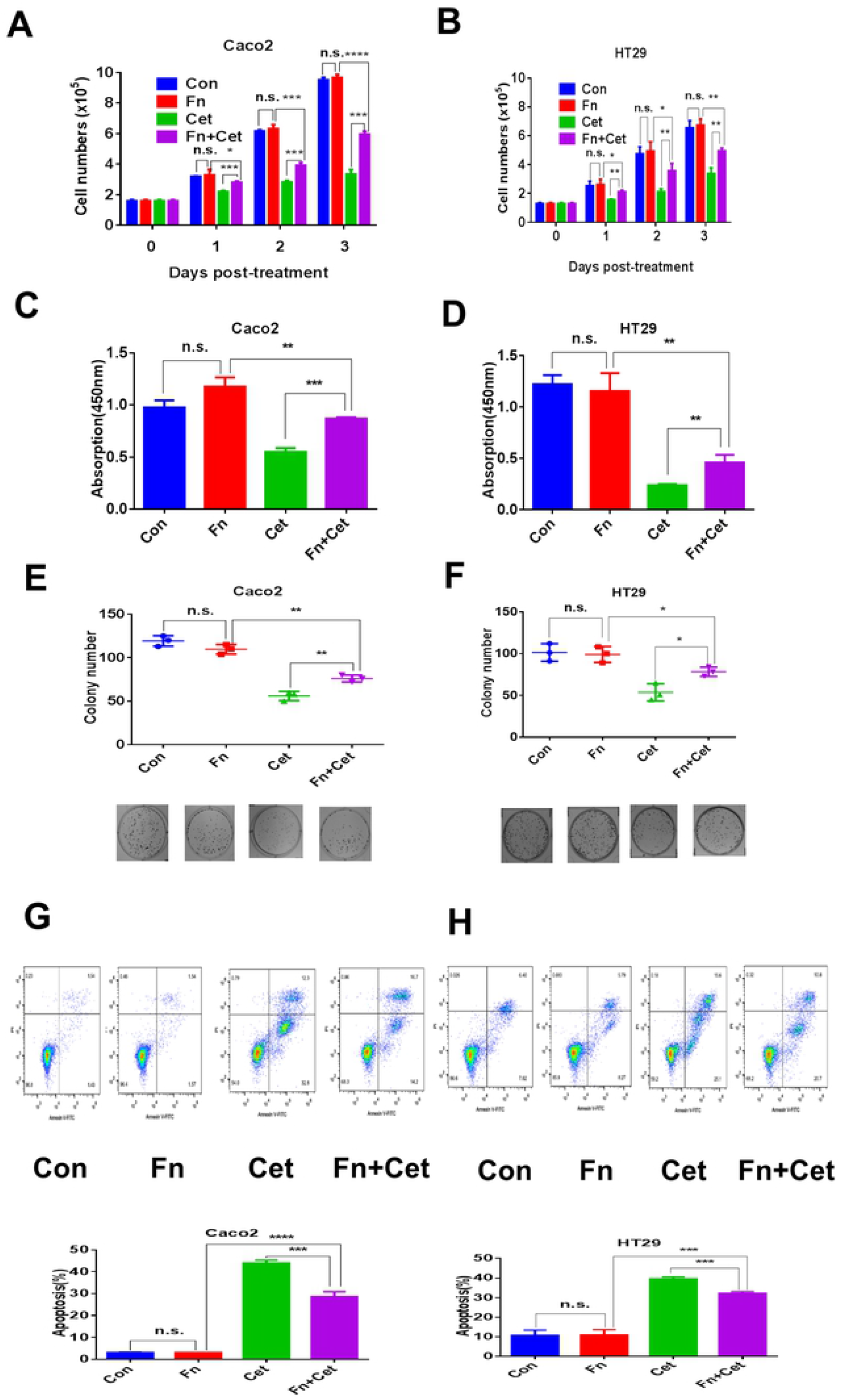
*Fn* induces cetuximab resistance in CRC cells *in vitro*. (A and B) Cell number was detected by a hemocytometer in Caco2 and HT29 cells with different conditions at indicated time. N = 3 independent experiments, unpaired Student *t* test. (C and D) Cell absorption was detected by spectrophotometer in Caco2 and HT29 cells with different conditions after 48h. N = 3 independent experiments, unpaired Student *t* test. (E and F) Caco2 and HT29 cells were cultured with different conditions and colonies were counted after 14 d. N = 3 independent experiments, unpaired Student *t* test. (G and H) Apoptosis was detected by flow cytometry in Caco2 and HT29 cells with different conditions after 48h. N = 3 independent experiments, unpaired Student *t* test. n.s., not significant, \**P* < 0.05, \*\**P* < 0.01, \*\*\**P* < 0.001, by unpaired Student *t* test.. Con, control; *Fn, Fusobacterium nucleatum* (MOI:100:1); Cet, cetuximab (100 μg/ml).

Next, we observed that *Fn* (MOI = 100:1) had no effect on the viability of Caco2 and HT29 cells than the untreated cells, but, it significantly reduced the effect of cetuximab on viability of these cells (Figures 2C *p*=0.0001 and 2D *p*=0.0065). Further, *Fn* (MOI = 100) did not affect the cellular transformation of Caco2 and HT29 cells than the untreated cells (Figure 2E *p*=0.1096 and 2F *p*=0.7899). While cetuximab reduced the cellular transformation in these cells, co-culturing them with *Fn* significantly reduced the effect induced by cetuximab (Figure 2E *p*=0.0064 and 2F *p*=0.0214). Furthermore, *Fn* (MOI = 100:1) had no effect on apoptosis in Caco2 and HT29 cells than the untreated cells. However, while cetuximab induced apoptosis in Caco2 and HT29 cells, co-culturing them with *Fn* significantly reduced the effects induced by cetuximab (Figure 2G *p*=0.0005 and 2H *p*=0.0005). Therefore, these results indicate that *Fn* inhibited the effects of cetuximab on cell viability, transformation, and apoptosis in Caco2 and HT29 cells.

Additionally, the Caco2 cells were inoculated into nude mice, and the mice were treated with cetuximab and/or *Fn*. Cetuximab significantly inhibited the tumor growth *in vivo*. While *Fn* alone had no effect on the growth of tumor cells, it inhibited the effect of cetuximab on tumor growth (Figure 3A–3E, Figure 3D *p* = 0.0044 and Figure 3E *p*=0.0008). Therefore, these results indicate that *Fn* confers resistance to reduction in tumor growth induced by cetuximab.

**Figure 3.**
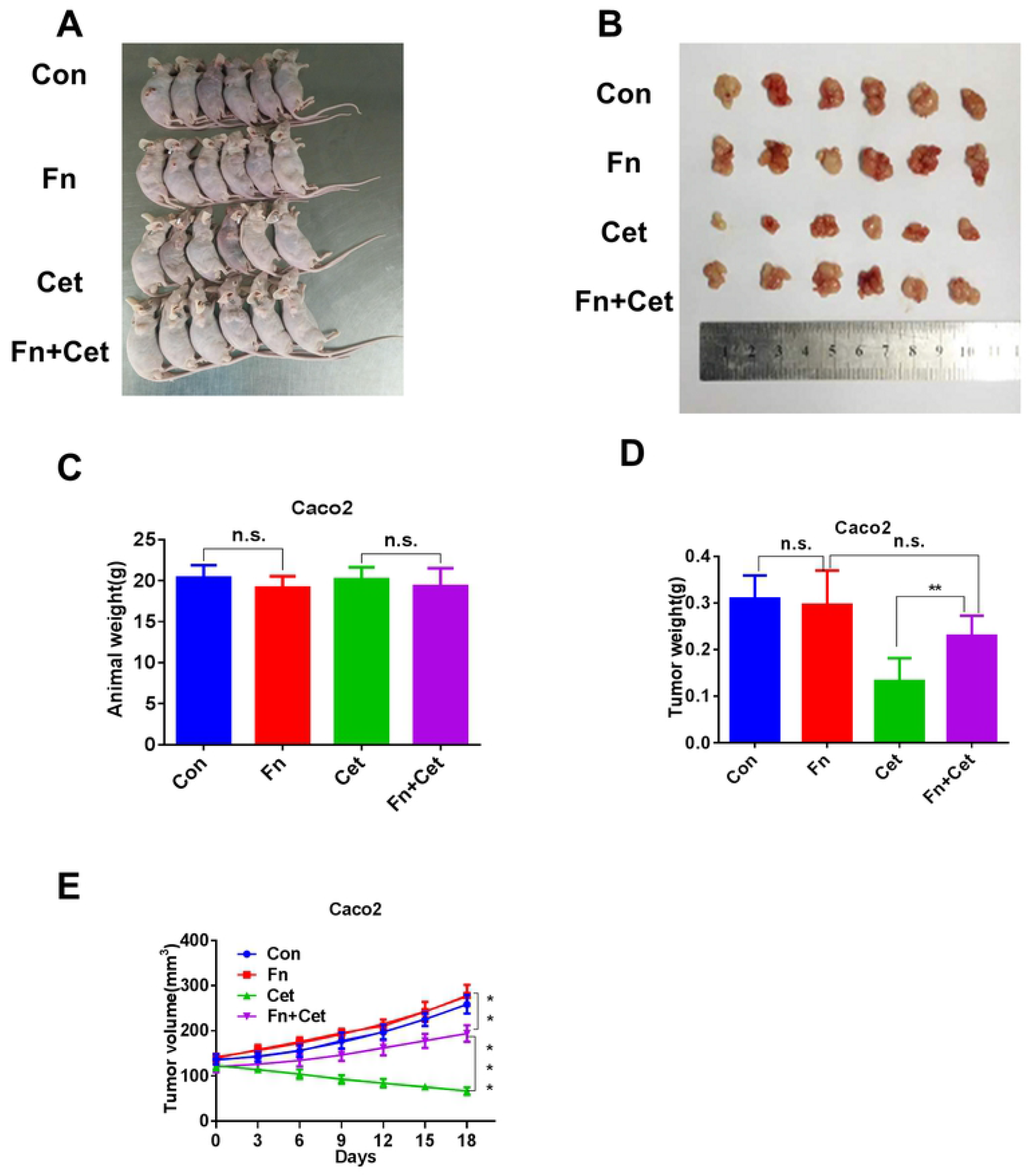
*Fn* induces cetuximab resistance *in vivo*. (A) Representative data of animal in nude mice bearing colorectal cancer cells in different groups. (B) Representative data of tumors in nude mice bearing colorectal cancer cells in different groups. (C) Statistical analysis of mouse weights in different groups, n = 6/group, unpaired Student *t* test. (D) Statistical analysis of tumor weights in different groups, n = 6/group, unpaired Student *t* test. (E) Statistical analysis of tumor volumes in different groups, n = 6/group, unpaired Student *t* test. Data represent mean ± s.d. \*\**P* < 0.01, \*\*\**P* < 0.001 by unpaired Student *t* test. n.s., not significant. Con, control; *Fn*, *Fusobacterium nucleatum*; Cet, cetuximab.

Taken together, the observations validate that *Fn* inhibits the effects of cetuximab on the CRC cell proliferation, viability, transformation, apoptosis, and tumor growth *in vivo*.

### *Fn* promotes activation of the PI3K/AKT and JAK/STAT3 signaling pathways

Next, we set to understand the mechanism of how *Fn* influences resistance to cetuximab. First, we found that *Fn* did not affect the cell cycle in Caco2 and HT29 cells after coculturing for 48 hours (Figure 4A and 4B). Further, since cetuximab and other mAbs bind the EGFR, blocking the activation of receptor tyrosine kinases and downstream signaling pathways associated with cell survival, proliferation, angiogenesis, and metastasis^4,50^, we checked whether EGFR or downstream signaling pathways in CRC cells were potential targets of *Fn*. Among the downstream signal pathways, the involvement of the RAS-RAF-MEK-ERK, PI3K-AKT, and JAK-STAT3 signaling pathways has been confirmed in conferring resistance to anti-EGFR mAbs^3,23,24^ Since deregulation of the PI3K/AKT and JAK/STAT3 pathways confers an anti-apoptotic and multidrug resistance phenotype to tumor cells^23,24^, we checked the effect of *Fn* on the activation of these pathways in CRC cells. Our western blot analysis indicated that the levels of P-AKT and P-STAT3 were increased in Caco2 (Figure 4C) and HT29 (Figure 4D) cells co-cultured with *Fn*. The levels of P-AKT and P-STAT3 were reduced in Caco2 (Figure 5A) and HT29 (Figure 5B) cells treated with cetuximab. The levels of P-AKT and P-STAT3 were still increased in Caco2 (Figure 5C) and HT29 (Figure 5D) cells treated with cetuximab followed infected with *Fn*. Thus, these results confirm the involvement of the PI3K/AKT and JAK/STAT3 pathways in conferring cetuximab resistance upon exposure of CRC cells to *Fn*.

**Figure 4.**
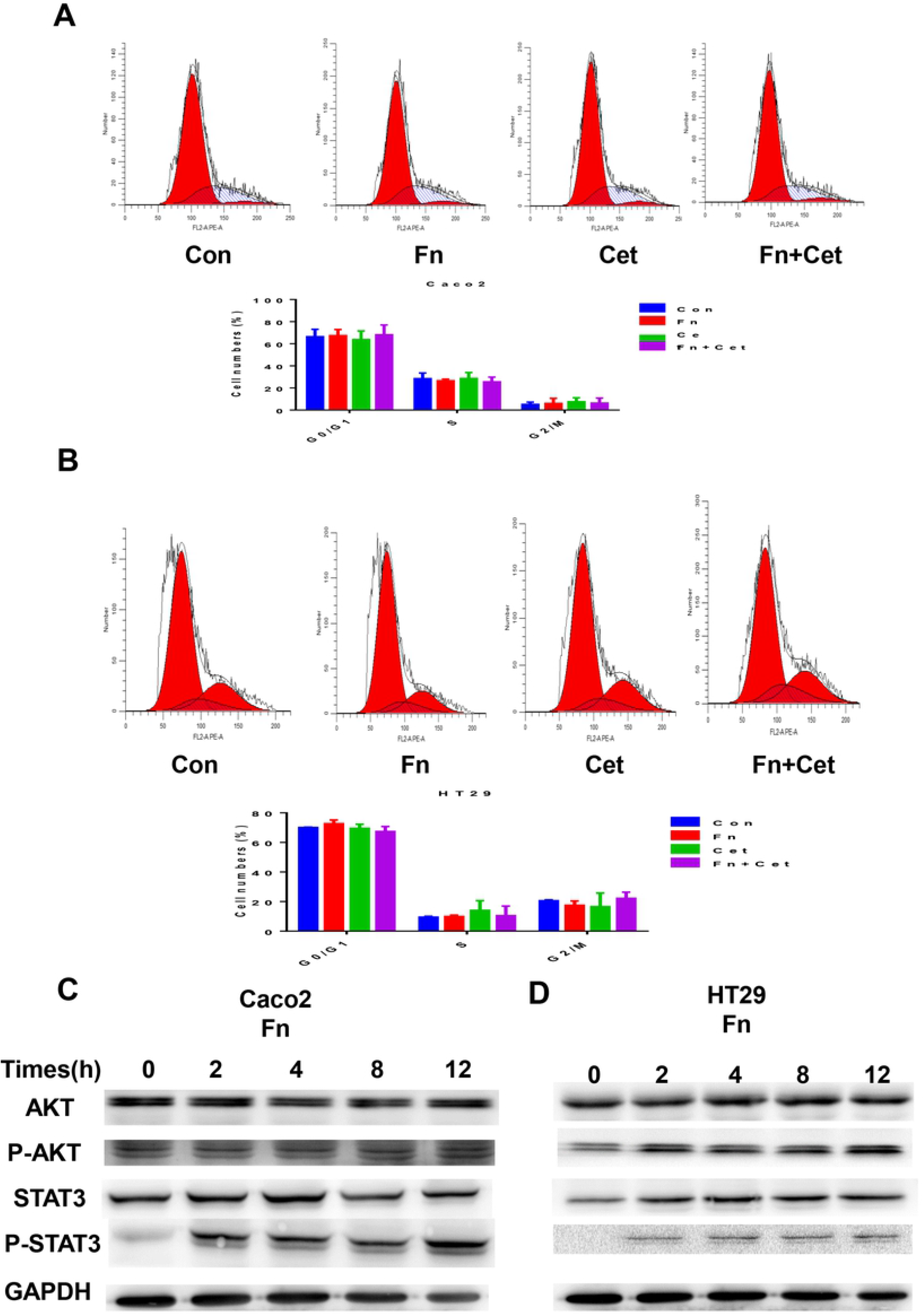
*Fn* promotes activation of the PI3K/AKT and JAK/STAT3 signaling pathways. (A) The cell cycle distribution of Caco2 cells with different treatment was determined by flow cytometry-based assay after 48 hours. N=3, unpaired Student *t* test. (B) The cell cycle distribution of HT29 cells with different treatment was determined by flow cytometry-based assay after 48 hours. N=3, unpaired Student *t* test. (C and D) Immunoblots of Caco2 and HT29 cell lysates from Caco2 and HT29 treated with *Fn* at indicated time. GAPDH served as the loading control. n.s., not significant. Con, control; *Fn*: *Fusobacterium nucleatum* (MOI: 100:1); Cet: cetuximab (100 μg/ml).

**Figure 5.**
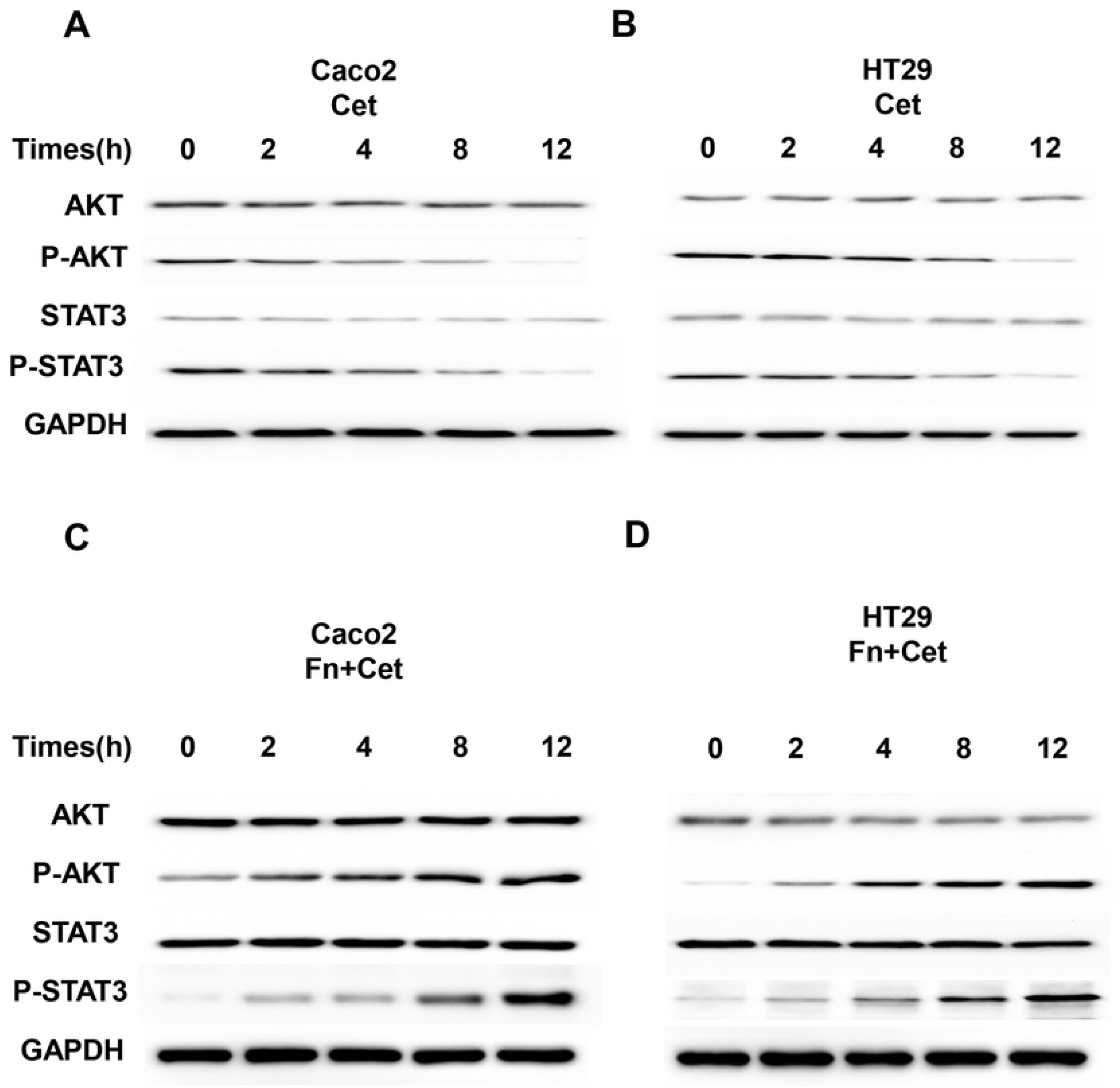
*Fn* promotes activation of the PI3K/AKT and JAK/STAT3 signaling pathways. (A) Immunoblots of Caco2 cell lysates from Caco2 treated with cetuximab at indicated time. GAPDH served as the loading control. (B) Immunoblots of HT29 cell lysates from HT29 treated with cetuximab at indicated time. GAPDH served as the loading control. (C) Immunoblots of Caco2 cell lysates from Caco2 treated with cetuximab followed infected with *Fn* at indicated time. GAPDH served as the loading control. (D) Immunoblots of HT29 cell lysates from HT29 treated with cetuximab followed infected with *Fn* at indicated time. GAPDH served as the loading control. *Fn*: *Fusobacterium nucleatum* (MOI: 100:1); Cet: cetuximab (100 μg/ml).

## Discussion

Studies suggest that mCRC is a genetic heterogeneous disease and that tumors appearing from diverse sites of the rectum have different clinical outcomes. The morbidity of left-sided and right-sided CRC varies, with about two-thirds of the patients presenting with the left-sided CRC^51^. Clinically, the right-sided CRC (RC) are more familiar in women, are diploid in nature, and more commonly associated with indicators of poor prognosis, such as mutations in the *RAS* and *BRAF*^52^, activation of MAPK signaling pathway, CpG island methylator phenotype-high, microsatellite instability, and mucinous histology^53,54^. Whereas, the left-sided CRC (LC) are common in men, show aneuploidy, chromosomal instability, HERI amplification, mutations in the *KRAS*, *DCC*, and *P53*, and gene expression profile consistent with the sensitivity of mAbs against the EGFR^9,55^. Furthermore, genetic non-polyploid CRC is frequent in the RC, whereas familial adenomatous polyposis develops in the LC^53,56^. Thus, patients with RC commonly associate with a poor prognoses than those with LC^57–59^, and they appear different during endoscopy^53^. Moreover, CRC shows genomic and epigenetic transformation via interactions between cancer cells, immune cells, and microorganisms that vary between RC and LC patients^60^.

Metagenomics and transcriptomics analyses have illustrated that the intestinal microorganisms, specifically *Fn*, participate in the progression of CRC^32,42,61^. Studies indicate that virulence factors of *Fn* associated with CRC, and it has been shown to invade the human epithelial cells, promote inflammation, and induce WNT signaling that contributes to tumor cell growth through FadA adhesion^43^. Another virulence factor, an autonomic transporter protein, Fap2, has been demonstrated to interact with the tight junction. This interaction protects tumor cells from host immune cell attack^44^. Some studies have demonstrated that the Fap2 induces interaction with Gal-GalNAc, a host polysaccharide overexpressed in CRC, so that it promotes adhesion and invasion of *Fn* in epithelial and endothelial cells^62^.

Here, we found *Fn* to promote resistance of CRC to cetuximab, *in vitro* and *in vivo*(Cetuximab was added and the results showed that it was present to increase apoptosis and reduced cell proliferation, viability and transformation even in cells cocultured with *Fn*. These results indicated that *Fn* was found to cause partial resistance to cetuximab.). The *Fn*-mediated activation of PI3K/AKT and JAK/STAT3 signaling pathways indicates the survival adaptation of cells regardless of inhibition of the EGFR. While our analysis indicates induction of P-AKT and P-STAT3 to promote cetuximab resistance in *Fn*-infected CRC cells, we cannot neglect the involvement of mechanisms other than the PI3K/AKT and JAK/STAT3 signaling pathways. For example, *Fn* can enhance cancer formation in the CRC cells via interaction with E-cadherin and direct activation of WNT signaling pathway^43^, which has been implicated in resistance to cetuximab^46^. Moreover, the detailed mechanism by which *Fn*-mediated activation of the PI3K/AKT and JAK/STAT3 signaling pathway that confers cetuximab resistance remains to be studied. Therefore, strategies need to be devised to unravel the crosstalk between cetuximab and *Fn*.

According to the 2019 NCCN guidelines, FOLFOX/CAPEOX/FOLFIRI/FOLFOXIRI ± bevacizumab has been recommended as first line therapy, while FOLFOX/FOLFIRI ± EGFR-targeted mAbs has been recommended as second-line therapy for patients with *RAS/BRAF* WT RC. Since previous report suggest that the patients with RC show abundance of *Fn* than in LC^38–40^, our study partly explains why *Fn*-high CRC patients present with poor prognoses^41,42^. Moreover, the study also attempts to describe why patients with *Fn*-high *RAS/BRAF* WT RC experience poor response to cetuximab than in patients with *RAS/BRAF* WT LC patients. Furthermore, our study raises an important clinical question that whether cetuximab is suitable for the treatment of CRC patients with high amounts of *Fn* irrespective of the site, RC or LC? Previous studies have shown that *Fn* inuduces CRC resistance to Oxaliplatin and 5-FU (FOLFOX) by Modulating Autophagy^42^. Although our results showed that *Fn* caused partial, not complete resistance to cetuximab, they still had some clinical significanc. Based on the analysis and previous studies, we recommend *Fn*-high patients to be treated with a combination of anti-EGFR mAb, anti-*Fn* therapy, and/or inhibitors of the PI3K/AKT and JAK/STAT3 pathways, Thus, it is significative to measure the amounts and target *Fn*, and its associated pathways, differentially based on the levels of *Fn* in the tumor.

Nevertheless, the relation between the intestinal microbiota and therapeutic outcome appears complicated, multifactorial, and of special context. Finally, we anticipate that the tumor microenvironment (TME)-targeted therapy should enhance the effects of anti-EGFR mAbs, and improve patient outcome.

## Contributions

HH, WQ, and YL performed the experiments, analyzed the data, and prepared the manuscript draft. HH, LW, and HY set up the experiments and repeated the key experiments. HH, BBS, XLY conceived the work, analyzed the data, and prepared the manuscript. All authors critically revised the manuscript, approved the final version, and agreed to be accountable for all aspects of the manuscript.

## Acknowledgments and funding

This work was supported by grants from the National Natural Science Foundation of China (grant numbers 81773360).

